# Sustained dynamics of saccadic inhibition and adaptive oculomotor responses during continuous exploration

**DOI:** 10.1101/2025.09.28.679009

**Authors:** Celeste Cafaro, Giovanni Cirillo, Giovanna Vermiglio, Carlo Cavaliere, Alessio Fracasso, Antimo Buonocore

**Author notes:** Co-first authorship (Celeste Cafaro, Giovanni Cirillo). Co-last authorship (Alessio Fracasso, Antimo Buonocore).

## Abstract

In natural environments, stimuli often recur across time and space, requiring the visual system to remain sensitive to novelty while managing predictability. A central question in systems neuroscience is how motor systems adapt to repeated sensory events without compromising responsiveness. We investigated this adaptive capacity using saccadic inhibition (SI)—a reflexive suppression of eye movements triggered by sudden visual transients—as a probe of oculomotor dynamics during naturalistic viewing. Human participants (N = 21) freely explored visual arrays while brief gaze-contingent flashes appeared five times at random intervals, either foveally or parafoveally. SI reliably occurred ∼120 ms post-flash across repetitions and locations, indicating robust sensory-driven inhibition. However, the rebound phase—reflecting saccade reprogramming—showed a progressive decline. In a second experiment (N = 19), only the first or the fifth flash was visible on each trial. In this case, neither inhibition nor rebound was altered, suggesting that the rebound decline is driven by repeated sensory stimulation rather than exploration time. This dissociation reveals selective habituation of motor re-engagement mechanisms, while reflexive inhibitory gating remains stable. We propose that inhibition is mediated by circuitry that transiently suppresses saccade initiation and resists habituation. By contrast, the weakening rebound reflects a separate, habituation-prone route that reduces saccade generation to irrelevant events. Functionally, this imbalance implies a recalibration within the saccade generator, preserving inhibitory capacity while constraining motor output. Our findings uncover a distinct form of oculomotor habituation and demonstrate how SI reveals dynamic decoupling of sensory input and motor output under repeated stimulation.

**Significance Statement:** The ability to interrupt and resume eye movements in response to environmental changes is fundamental to visual exploration. We investigated how this process unfolds across repeated visual transients in naturalistic conditions. Our findings show that saccadic inhibition remains stable across time, whereas the subsequent rebound phase habituates. This dissociation suggests that distinct processes mediate reflexive interruption and motor recovery habituation. Our data indicate that dynamic modulation of motor gating circuits is a mechanism for optimizing oculomotor behavior. These results deepen our understanding of visual-motor coordination, provide insight about the constraints governing the underlying system circuitry, and may inform clinical tools for assessing sensorimotor adaptability in health and disease.

## Introduction

Exploring the environment is essential for survival, requiring the nervous system to continuously integrate dynamic sensory input with motor plans (Hayhoe, 2017). Vision plays a dominant role in this process, and eye movements actively sample the visual world, redirecting the gaze toward relevant stimuli. While these movements are often goal-directed, they are also shaped by sudden external events. The visual system must therefore remain responsive to novelty while adapting to environmental regularities (Pascucci et al., 2019). Within this perception–action loop (Fuster, 2004), repeated stimulation is common. In response, the brain might engage sensory adaptation—a largely automatic reduction in neural responsiveness to persistent or fast recurring inputs—that enhances coding efficiency and aligns perception with context (Webster, 2011; Solomon and Kohn, 2014).

Adaptation occurs across various visual features, including contrast, orientation, motion, and complex forms like faces (Sekuler and Ganz, 1963; Blakemore and Campbell, 1969; Georgeson and Harris, 1984; Greenlee and Heitger, 1988; Anstis et al., 1998; Webster and MacLeod, 2011). From a neural perspective, adaptation reduces firing in repeatedly stimulated circuits, from the retina to higher-order areas (Smirnakis et al., 1997; Kohn, 2007; Benucci et al., 2013; Torrado Pacheco et al., 2019). Oculomotor areas also adapt: the superior colliculus (SC) shows attenuated responses to repeated transients (Boehnke et al., 2011; Basso and May, 2017), and the frontal eye fields (FEF) exhibit strong adaptation to sequential flashes (Bruce and Goldberg, 1985; Mayo and Sommer, 2008).

External events can also recur over longer time scales. Whereas fast adaptation reflects transient sensory adjustments, habituation denotes a slower, learned reduction in behavioral responses to repeated, irrelevant stimuli (Boehnke et al., 2011). Unlike adaptation, habituation engages higher-order modulation shaped by attention, expectation, and context, thereby supporting goal-directed behavior in environments rich with distractors (Burleson et al., 2025). A central question in systems neuroscience is how motor circuits respond to repeated sensory stimulation, balancing responses that adapt or habituate with the need to react rapidly and flexibly to external stimuli.

Saccadic inhibition (SI) provides a paradigm to probe visuomotor control: an abrupt visual transient suppresses eye movements ∼90 ms later, followed by a rebound at 120–150 ms reflecting motor reprogramming (Reingold and Stampe, 1999, 2002, 2004; Buonocore and McIntosh, 2008; Edelman and Xu, 2009; Bompas and Sumner, 2011). This inhibitory–rebound cycle is shaped by stimulus and task factors (e.g., location, salience, attention, content) (Bompas and Sumner, 2011; Buonocore and McIntosh, 2012, 2013; Buonocore et al., 2017a; Taylor et al., 2024) and extends beyond saccades to microsaccades, smooth pursuit, and even fixation drift (Rolfs et al., 2008; Kerzel et al., 2010; Buonocore et al., 2017b; Buonocore et al., 2019; Malevich et al., 2020). Together, these findings identify reflex-like inhibition as a unifying mechanism at the core of oculomotor control.

Neurally, SI is thought to act downstream of the SC, potentially involving omnipause neurons (OPNs) that gate saccade initiation (Buonocore and Hafed, 2023). Subcortical structures such as the SC, as well as cortical interventions targeting the FEF, LIP, and other motor areas, might support the reinstatement of eye movements. While SI has been well studied in isolated trials, its evolution under repeated stimulation—as it might occur during natural viewing—remains poorly understood. Evidence indicates that the visual component driving inhibition adapts to rapidly presented visual stimuli, resulting in reduced effects (Shan and Edelman, 2023). However, it is not yet established how the oculomotor system responds to repeated flashes over longer time scales that more closely approximate natural viewing conditions.

We hypothesize that SI reflects two distinct mechanisms: a fast, visually driven inhibition phase subject to short-timescale adaptation (Shan and Edelman, 2023), and a motor-driven rebound phase vulnerable to slower habituation. Rapid successive transients may reduce inhibition through sensory adaptation, whereas at longer intervals (∼1 s), inhibition remains stable while the motor system disengages from post-inhibition recovery. In this study, participants freely explored visual scenes while gaze-contingent transients appeared foveally or parafoveally. If inhibition and rebound were tightly coupled, both would decline together, implying a single governing process. Instead, evidence for their dissociation would support independent mechanisms: sensory processes regulating inhibitory strength and motor processes regulating recovery dynamics, enabling selective tuning of responsiveness without uniformly suppressing downstream motor output. By dissociating these components of oculomotor behavior, our findings advance understanding of how the brain balances responsiveness with efficiency under repeated interference.

## Materials and methods

### Participants

Experiments 1 and 2 were eye-tracking experiments on human participants. Experiment 1 included a sample of twenty-one participants (17 females and 4 males), while Experiment 2 included a sample of nineteen participants (14 females and 5 males). All participants had normal or corrected-to-normal vision, reported no neurological or visual impairments, and were naïve to the purpose of the experiment. The sample comprised university students from the Department of Educational, Psychological and Communication Sciences from the Psychology Degree Course of Suor Orsola Benincasa University. Written informed consent was obtained from all participants in compliance with the 2024 Declaration of Helsinki. The study received ethical approval from the local ethics committee of Suor Orsola Benincasa University, Naples.

### Apparatus and stimuli

In both Experiment 1 and 2, participants were seated with their eyes aligned to the center of the display at a viewing distance of 65 cm, with their heads stabilized using a chin and forehead rest, to avoid head movements and ensure that the eyes were positioned approximately in front of the center of the screen. Experimental sessions were conducted in a dark room, illuminated only by the experimental display monitor and the operator’s monitor, which was positioned behind the participant and facing away from them. Eye movements were recorded using a LiveTrack Lightning eye tracker (Cambridge Research Systems), sampling both eyes at 500 Hz and determining the direction of gaze from the pupil and Purkinje centers. Before the start of the experiment, a binocular calibration was performed using nine locations arranged in a three-by-three grid. The calibration procedure was performed with the LiveTrack Viewer software, and calibrations were accepted if the error for each eye was below 0.5 degrees of visual angle. Both eyes were then tracked throughout the entire experiment. The experimental protocol and stimuli were generated using MATLAB R2023a (MathWorks, Inc.) in combination with Psychophysics Toolbox 3.0.14 (Brainard, 1997; Pelli, 1997). All stimuli were presented in black on a mid-gray background on a 27-inch LCD monitor with a resolution of 1920 by 1080 pixels and a refresh rate of 144 Hz (6.9 ms). A small blue fixation dot, with a radius of about 0.30 degrees (10 pixels) served as a fixation point. Stimuli consisted of screens displaying randomly generated geometric figures. Depending on the task (see Procedure), each screen contained either horizontal/vertical lines or circles/squares presented in completely random positions. All the geometric shapes were about two degrees in size. The number of elements was randomly determined on each trial between 19 and 29. In particular, one display might contain more horizontal than vertical lines (or vice versa), with a difference of 1 to 3 elements. The same logic applies to circles and squares. During the stimulus presentation, a black circle two degrees in size appeared at various retinal locations, either within the fovea (Experiment 1 and 2) or 5.6 degrees in the periphery (only Experiment 1), always on the left side of the fovea. Manual responses to the shape-number estimation task were collected using a five-button USB response pad from The Black Box Toolkit.

### Procedure

Both experiments consisted of four blocks, each comprising two tasks: lines (Task one) or shape (Task two) number estimation. At the beginning of each block, a brief on-screen sentence informed participants whether the upcoming comparison of the number of elements would involve lines or shapes. In each task, participants were required to explore the visual display and judge the numerosity of geometric items (Figure 1). Specifically, participants were instructed to scan the entire screen to estimate whether the display contained more vertical or horizontal lines (Task one), or more circles than squares (Task two). The sequence of tasks was randomized across blocks for each participant. Participants initiated each task by pressing the central button on a five-button response box. On each trial, they were first asked to fixate on a blue dot presented at the center of the screen. After a variable delay between 1 and 1.5 seconds from fixation onset, a stimulus array was displayed for approximately 12 seconds. Following stimulus onset, participants freely explored the display for approximately 12 seconds. No fixation window was enforced during this phase, and no online fixation constraint was applied.

**Figure 1.**
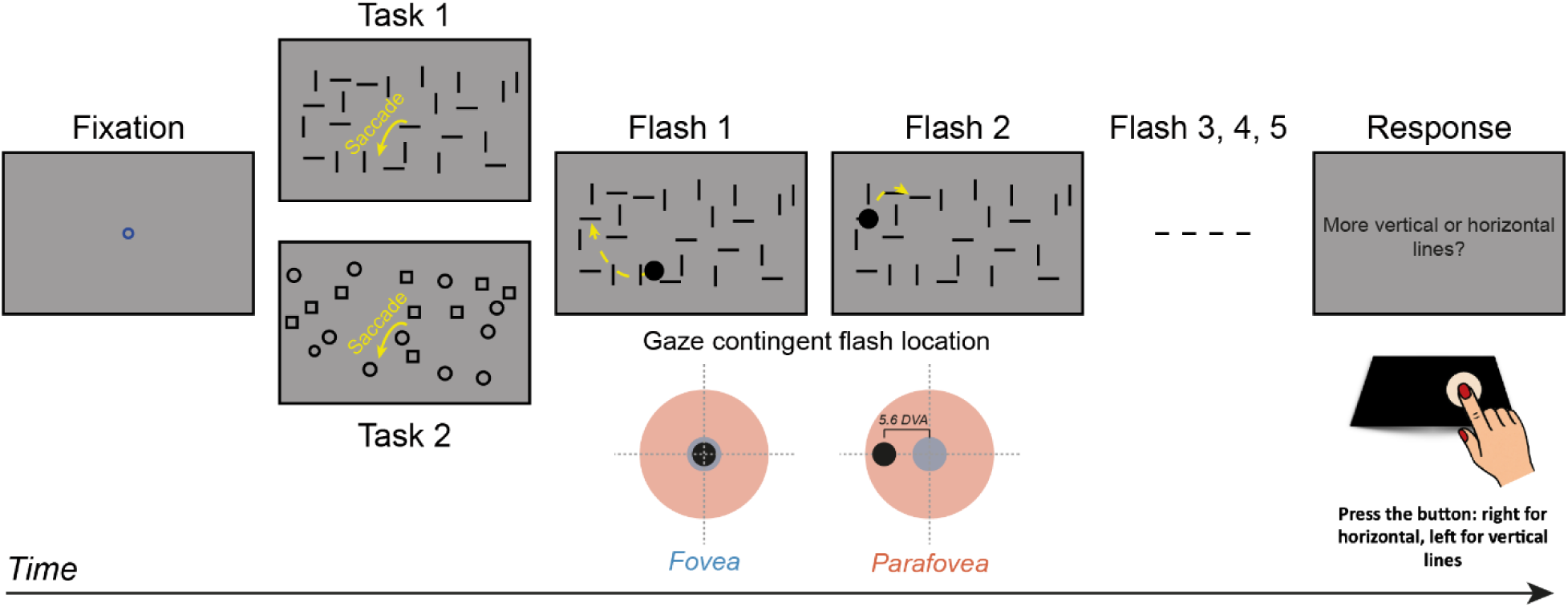
Procedure and stimuli used in the experiment. Participants fixated on a central dot for a variable duration (1 to1.5 s). A display containing geometric shapes, either lines (Task 1) or circles/squares (Task 2), was then presented for about 12 s. During visual exploration, a gaze-contingent black circle flashed for 80 ms either five times per trial (Experiment 1) or was visible only on the first or fifth presentations (Experiment 2). At the end of each trial, participants reported the number of figures observed using a button box. In Experiment 1, the transient stimulus could appear either at the fovea (blue) or 5.6 degrees of visual angle along the horizontal meridian in the parafovea (red), to the left of fixation. In Experiment 2, the transient was always presented at the fovea.

During the visual exploration phase, brief transient circles were presented for 80 ms, contingent on gaze position. Eye position was sampled at 500 Hz, and the display was refreshed at 144 Hz. During each refresh cycle, all available gaze samples were averaged to estimate the instantaneous eye position for stimulus updating. The expected gaze-contingent system delay (from gaze sampling to stimulus onset on screen), determined by sampling and refresh timing, was approximately 7ms. Given that saccadic inhibition effects were analyzed for saccades initiated ≥100 ms after flash onset, and that the flash duration was 80 ms (11 frames), this latency does not affect the interpretation of the reported inhibition dynamics. In Experiment 1, each trial included five such transients, with an inter-stimulus interval (ISI) ranging from 0.7 to 1.2 seconds. Depending on the condition, the transient circle was presented either at the fovea or at a fixed eccentricity along the horizontal meridian, about 5.6 degrees away from the fovea (see Apparatus and stimuli). In Experiment 2, the transient circle appeared as either the first or the fifth stimulus, depending on the experimental condition. The remaining four circles were flagged as “invisible”, meaning that a stimulus trigger was only sent for time alignment (See Experimental design and statistical analysis).

In both experiments, following the stimulus presentation, a response screen appeared. Participants were asked to indicate whether they had seen more circles or more squares, or more horizontal than vertical lines, using two designated buttons on the response box. The left button was used to report more circles or vertical lines, while the right button was used to report more squares or horizontal lines. No time limit was imposed for responses. After a response was given, the next trial began automatically. The experiments comprised 120 trials, organized into four blocks of 30 trials each, with 60 trials dedicated to each geometric task. In Experiment 1, across the entire experiment, 600 flashes were presented—300 at foveal and 300 at parafoveal locations—with 60 flashes delivered at each of five flash orders. In Experiment 2, a total of 120 visible flashes were presented, 60 flashes presented as first and 60 flashes presented as fifth. The remaining 480 flashes were classified as “invisible”. A break was provided after the first two blocks to allow participants to rest their eyes. The full session lasted approximately 30 minutes, including the break. Participants resumed the experiment by pressing the central button on the response box. At the end of the session, all participants were debriefed about the purpose of the study.

### Data preprocessing

For both experiments, continuous raw eye position data (recorded in Fick coordinates: longitude and azimuth) were segmented into trials and stored as matrices. Eye velocity was computed using a custom band-limited differentiator that combines smoothing and numerical differentiation using a symmetric kernel. This procedure follows the saccade detection pipeline described in Buonocore et al. (2017b), based on the methodological framework of Chen and Hafed (2013) and Hafed et al. (2009). Horizontal and vertical eye position signals, sampled at 1 kHz, were differentiated to obtain velocity components, from which radial eye velocity was computed. Radial acceleration was then obtained as the temporal derivative of radial velocity. Saccades were detected automatically using a combination of velocity and acceleration criteria. Specifically, radial eye velocity exceeding 30°/s and radial acceleration exceeding 900°/s² were used as thresholds for candidate events. Movement onset and offset were refined by iteratively tracing backward and forward in time until acceleration fell below threshold, allowing precise estimation of movement boundaries. Each trial was manually inspected to correct false positives and missed detections. Blinks, identified by abrupt discontinuities in eye position, were manually marked and excluded from analysis. Saccade amplitude was calculated as the Euclidean distance between eye position at movement onset and offset, and peak velocity was defined as the maximum value of the radial velocity trace.

### Experimental design and statistical analysis

To examine how saccadic responses evolved across repeated visual transients, we computed saccadic inhibition profiles separately for each flash location (foveal vs. parafoveal) and flash order (flash 1–5). By saccadic inhibition profile, we refer to the change in saccade rate (expressed in saccades per second, Hz) over time relative to flash onset (Reingold & Stampe, 2002; Buonocore & McIntosh, 2008). Each trial contained five flashes delivered at pseudo-random time points during free viewing. Consecutive flashes were separated by a variable inter-stimulus interval (ISI) constrained between 700 and 1200 ms. This temporal spacing ensured sufficient separation between successive flash-evoked responses. For the analysis, all saccades within each trial were extracted and aligned to flash onset by subtracting the time of each flash from the time of each saccade. For every flash, we defined a temporal epoch from −200 ms to +500 ms relative to flash onset. Because the minimum ISI exceeded the epoch duration, epochs corresponding to different flashes within the same trial did not overlap. Each trial, therefore, contributed five independent flash-locked epochs, labeled by flash order (1–5). These epochs were then pooled across trials and participants and analyzed separately by flash order and flash location. Saccades with a peak velocity exceeding 600 degrees per second (Experiment 1: 0.37%; Experiment 2: 0.38%) and those with an amplitude greater than 20 degrees of visual angle (Experiment 1: 2.45%; Experiment 2: 2.42%) were excluded from the rest of the analysis. Next, for each trial, we constructed histograms of saccade frequency within a time window spanning from 200 milliseconds before to 600 milliseconds after each flash onset, using a bin width of 10 milliseconds. These histograms were then converted into saccade rates per second by dividing the counts by the bin width and multiplying by 1000. The resulting histograms were grouped by participant and condition, distinguishing between foveal and parafoveal flashes (Experiment 1) and between flashes 1 through 5 (Experiments 1 and 2). In Experiment 2, we also constructed histograms for “invisible” flashes by aligning the saccades generated during visual exploration with the timestamps of the so-called “invisible” flashes, which were not drawn on screen. This procedure allowed for the visualization of the flat distribution of saccades aligned with the trigger event. For each participant and condition, the histograms were averaged and smoothed using the MATLAB function “smoothdata” with the parameter “movemean” and a window of seven samples. This smoothing procedure corresponds to a moving average over 70 milliseconds, centered on the fourth sample, and advanced in 10-millisecond steps. Edge effects were handled automatically by the function, which initiates full smoothing from the fourth sample onward. At the end of preprocessing, each participant contributed ten smoothed histograms—one for each flash-location and flash order condition in Experiment 1, and for flash visibility and flash order condition in Experiment 2. The averaged histograms were used to extract key temporal and dynamic features of the saccadic response.

To quantitatively assess the dynamics of saccadic inhibition, we extracted a set of descriptive parameters, means, and standard deviations, from the saccade histograms of each participant and condition during critical time periods of 50 ms durations. These parameters captured both the overall magnitude of the response and the timing of key features in the inhibition profile. In Experiment 1, we first measured the baseline saccade frequency (i.e. *Baseline frequency*), defined as the average rate of saccades in the pre-flash interval from −100 to −50 ms, which served as a reference for ongoing oculomotor activity before the onset of the transient. Following the flash, we identified the minimum saccade frequency (i.e., *Inhibition frequency*) within a predefined window (90 to 140 ms), reflecting the depth of the inhibitory response, as well as the latency to this minimum (i.e., *Inhibition latency*), indexing the time point at which inhibition reached its peak. We then turned to the rebound phase, capturing the transient increase in saccade frequency that typically follows inhibition (i.e. *Rebound frequency*). Here, we measured the maximum rebound frequency within a post-inhibition window (200 to 250 ms). In Experiment 2, all the parameters were the same, but we observed a small shift of about 20 ms in the saccadic inhibition profile. For this reason, we computed inhibition and rebound frequencies: the former between 110 and 160 ms, and the latter between 240 and 290 ms after flash onset. All these temporal windows are visually indicated by shaded rectangles in Figures 4 and 5, providing a structured summary of the key stages of the saccadic inhibition response, alongside a table listing all measures extracted in Experiments 1, and 2.

Each of these dependent variables (*Inhibition frequency, Inhibition latency, Rebound frequency*) was analyzed using linear mixed-effects models (LME). In Experiment 1, fixed effects included *Flash location* (foveal versus parafoveal), *Flash order* (from the first to the fifth flash), their interaction, and the pre-flash *Baseline frequency* as a continuous covariate. A random intercept for each Participant was included to account for inter-individual variability. For the dependent variables *Inhibition frequency*, *Inhibition latency,* and *Rebound frequency,* the general model structure was as follows, in Wilkinson notation (Wilkinson and Rogers, 1973):

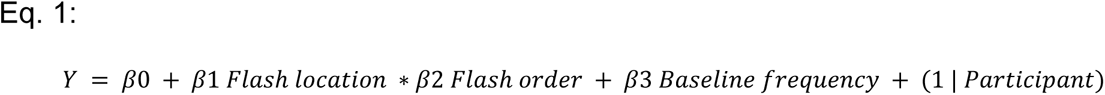

To examine whether rebound dynamics were explained by prior inhibition, we tested *Rebound frequency* as the dependent variable with a second model, extending the model in Eq.1 by adding *Inhibition frequency* as a continuous predictor (Eq. 2):

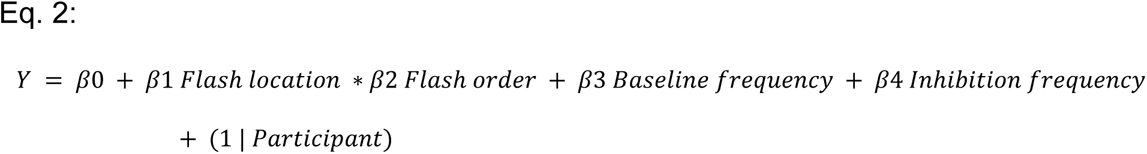

In Experiment 2, we compared *Rebound frequency* between the first and fifth flash using a paired-samples t-test. Additionally, we compared *Rebound frequency* for the fifth flash between Experiment 1 and Experiment 2 using an independent-samples t-test.

## Results

### Saccadic inhibition remains consistent across saccades of varying amplitudes, directions, and velocities

We first characterized eye-movement behavior qualitatively by analyzing standard saccade parameters across all trials (Figure 2). In Experiment 1, a total of 31,559 saccades were retained after manual screening to exclude detection errors (see Data preprocessing). As expected in free-viewing tasks (Foulsham et al., 2008; Foulsham and Kingstone, 2010), eye movements were predominantly horizontal (Figure 2A). Saccade amplitudes had a mode at approximately 2.5 degrees of visual angle (Figure 2B), consistent with serial object inspection. The main sequence (Figure 2C) exhibited the typical nonlinear relationship between saccade amplitude and peak velocity (Bahill et al., 1975; Buonocore et al., 2017b).

**Figure 2.**
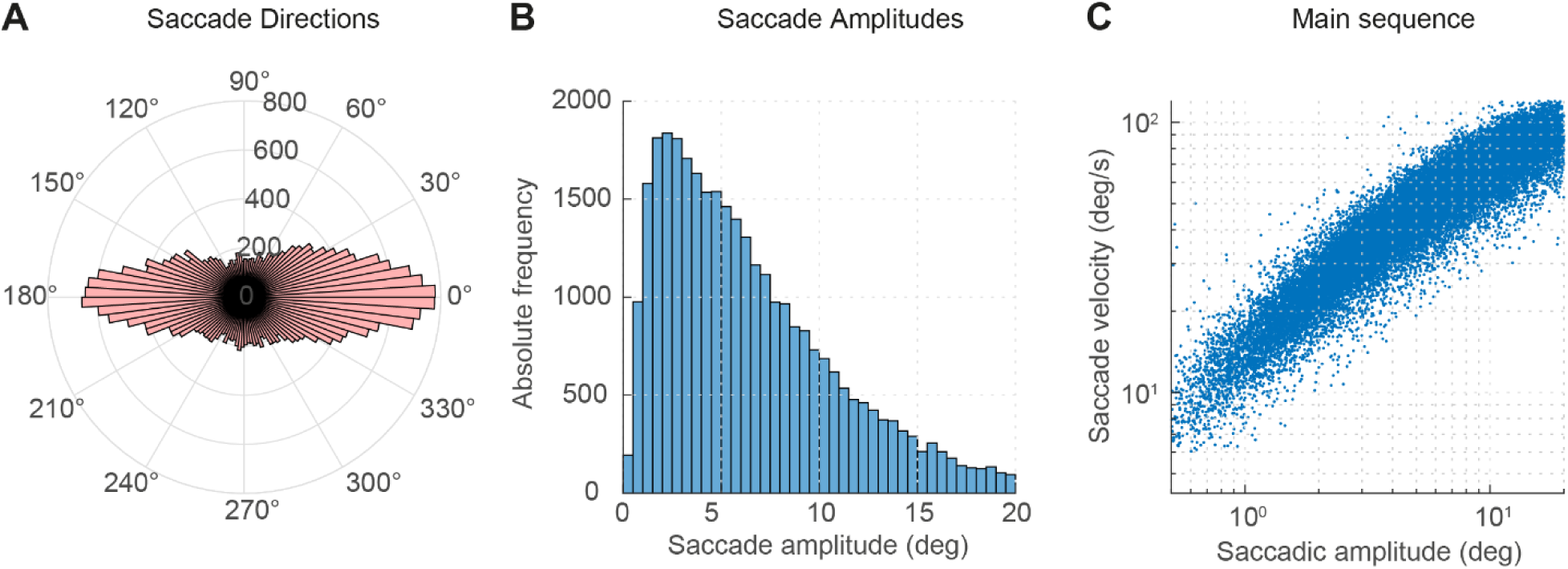
Global oculomotor behavior during free viewing (Experiment 1). (A) The distribution of saccade directions shows a predominant horizontal bias in visual exploration, as commonly observed in free-viewing tasks. (B) The distribution of saccade amplitudes is highly positively skewed, with a mode at ∼2.5 degrees of visual angle, consistent with serial object inspection strategies that favor smaller saccades. (C) From the main sequence, it is evident that peak velocity scales nonlinearly with amplitude, reflecting the canonical relationship. Note that the main sequence is plotted in log-log coordinates.

Temporal characteristics, including saccade and fixation durations (Figure 3A), were also consistent with established profiles of unconstrained visual exploration. Critically, despite substantial variability in saccade properties highlighted in Figure 2, a consistent pattern of saccadic inhibition emerged when saccades were time-locked to flash onset. A robust suppression of saccade frequency was observed approximately 120 ms post-flash across all participants (Figure 3B), confirming the reliability of SI under more viewing conditions. These results support the notion that saccadic inhibition is a stable and generalizable feature of oculomotor control, largely invariant to fluctuations in saccade amplitude, direction, or velocity (Reingold and Stampe, 2000, 2004).

**Figure 3.**
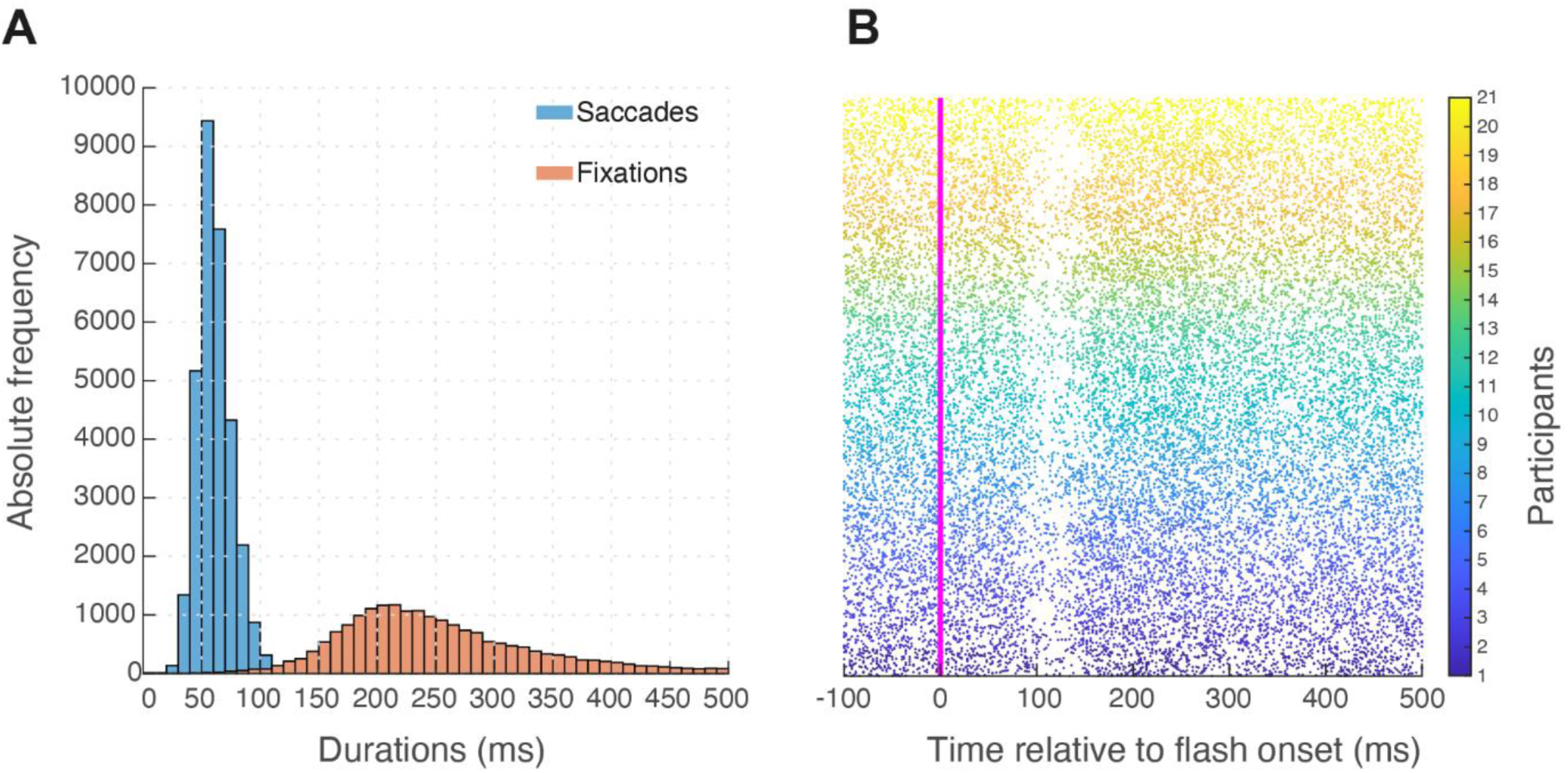
Distributions of saccade and fixation durations. A) Saccade and fixation durations fell within the expected range for unconstrained exploration. (B) When time-locked to flash onset (magenta vertical bar), saccades showed a robust suppression at ∼120 ms, confirming reliable saccadic inhibition during free viewing. Each dot represents a single saccade. The dataset includes 31,559 saccades after manual screening, with colors indicating individual subjects.

### Foveal flashes elicit stronger inhibition than parafoveal ones

We next investigated how the spatial location of visual transients modulates the temporal dynamics of saccadic inhibition and rebound. Specifically, we compared the effects of foveal and parafoveal flashes by analyzing their associated inhibition profiles (see *Experimental Design and Statistical Analysis*). Both flash types induced a pronounced inhibition peaking around 120 ms after onset (Figure 4A and B), with gray lines indicating consistent effects across individual participants. At visual inspection, foveal flashes (blue line, Figure 4A) produced stronger and more temporally precise inhibition compared to parafoveal flashes (red line, Figure 4B, see statistics below). This was followed by a sharper and more pronounced rebound in saccade frequency, mirroring the inhibition phase and again favoring foveal stimulation.

**Figure 4.**
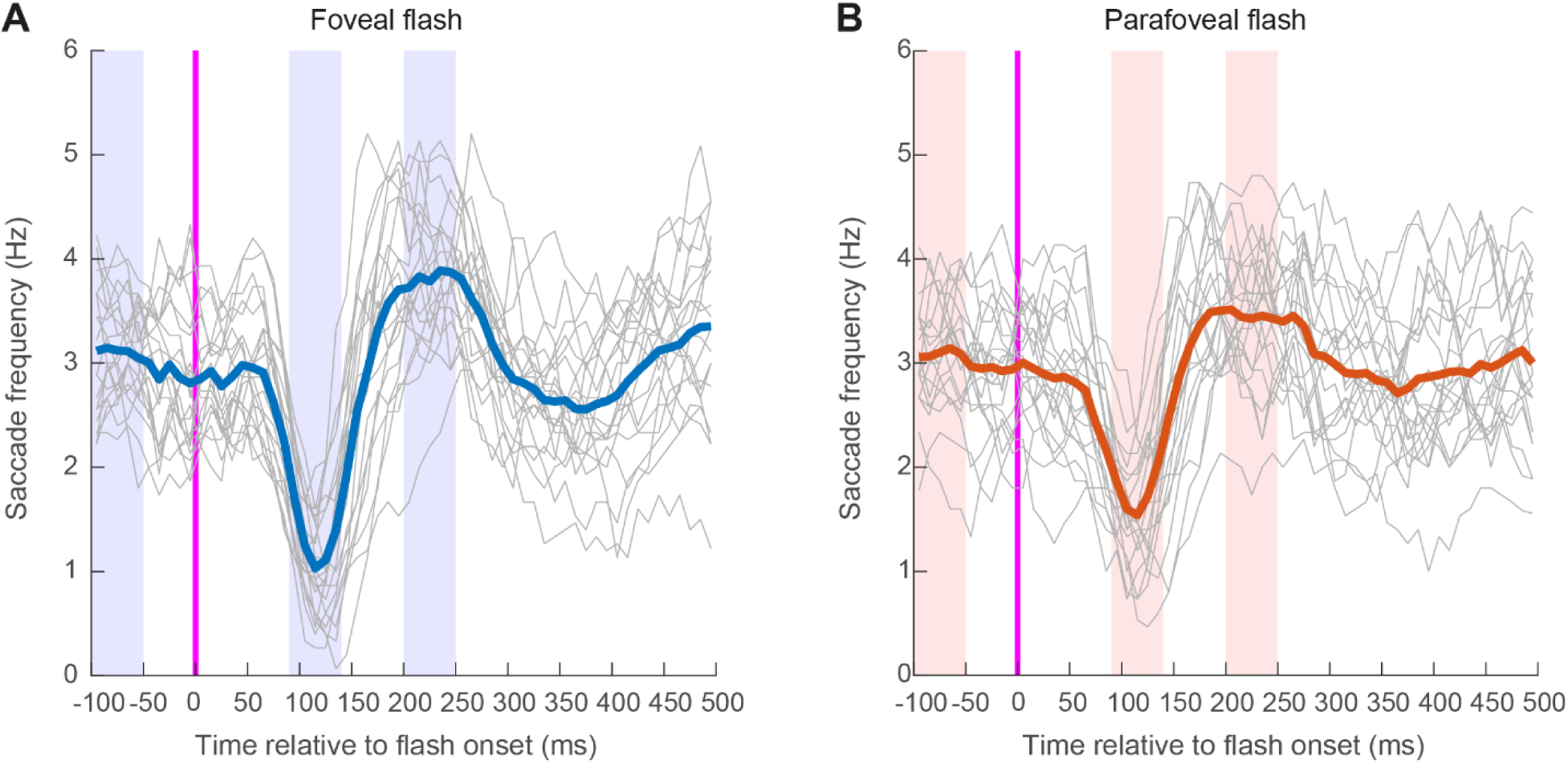
Saccadic inhibition profiles in foveal and parafoveal flashes (Experiment 1) Saccadic inhibition profiles for foveal (panel A, blue) and parafoveal (panel B, red) flashes showed a robust reduction in saccade generation ∼120 ms after onset. Foveal flashes produced stronger and more temporally precise suppression, followed by a sharper rebound in saccade rate ∼100 ms later. Gray lines represent individual participants, demonstrating consistent effects across subjects. Shaded blue and red rectangles indicate the time windows used to compute group averages for baseline, inhibition, rebound, and post-rebound measures (see *Experimental Design and Statistical Analysis* and Figure 2).

In Figures 5A and 5 B, we further partition inhibition profiles across five successive flashes (dark to brown lines indicate the first to fifth flash). Notably, the magnitude and latency of saccadic inhibition remained remarkably stable across repetitions, regardless of flash location. This suggests that while spatial position modulates the strength of the inhibitory response, the temporal dynamics of the mechanism are preserved across repeated stimulation. In contrast, a clear pattern emerged in the rebound: later flashes increasingly hindered the generation of new saccades, weakening the rebound phase of the saccadic inhibition profile. In addition, baseline levels appeared to be affected by flash repetition, with later saccades starting from a reduced frequency. In the next sections, we provide statistical modeling of the saccadic inhibition parameters described in the *Experimental design and statistical analysis* section.

**Figure 5.**
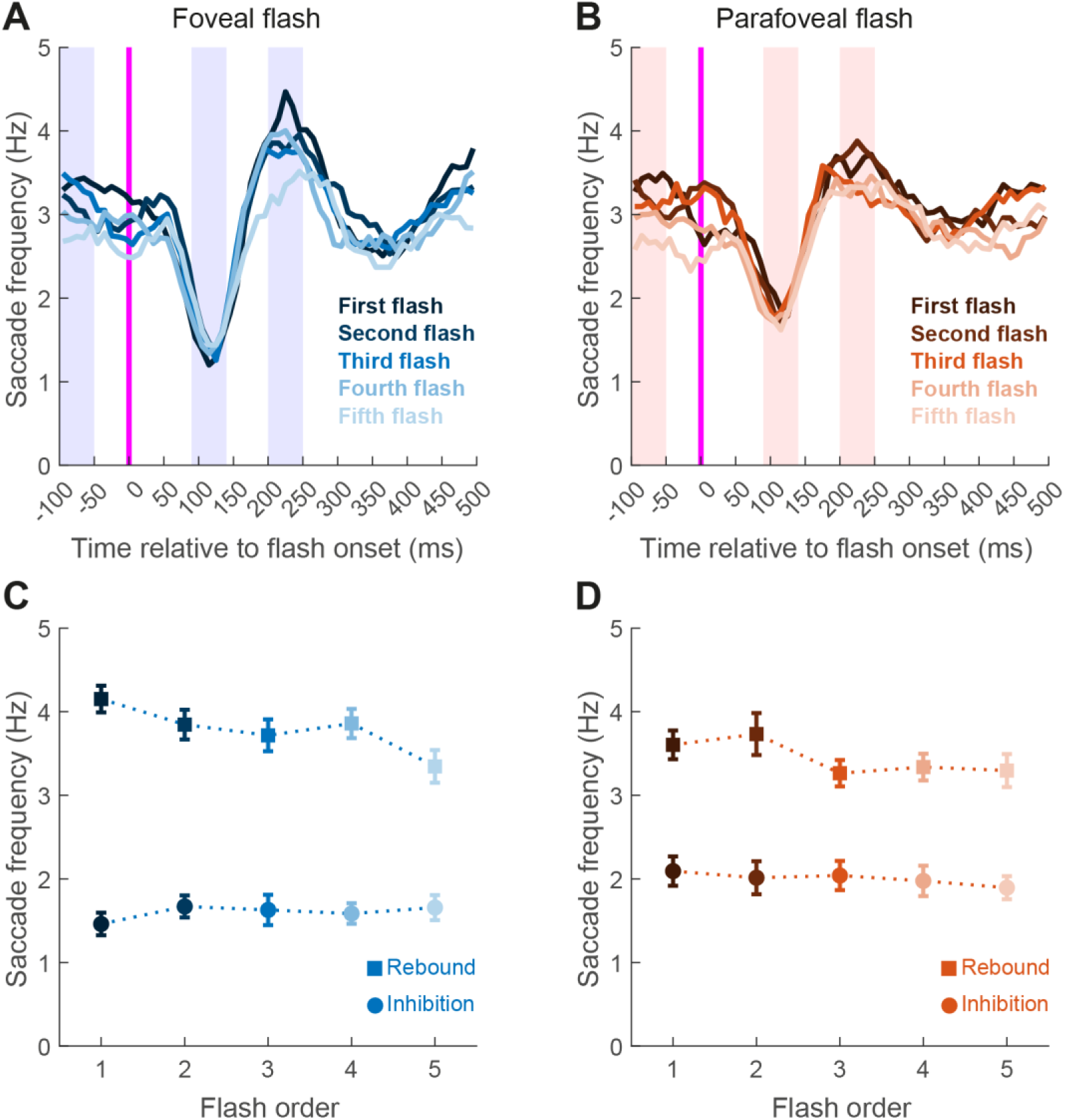
Saccadic inhibition profiles and parameters across flash order (Experiment 1). Saccadic inhibition profiles across flash order (1–5) showed a robust reduction in saccade generation ∼120 ms after onset. (A) Foveal flashes elicited stronger and more temporally precise suppression. The rebound in saccade rate occurred ∼100 ms later, being clearly marked for the first flash but progressively less pronounced for subsequent flashes. (B) Parafoveal flashes followed the same pattern, though with an overall weaker rebound. Shaded rectangles indicate the time windows used to compute group averages for baseline, inhibition, rebound, and post-rebound measures (see *Experimental Design and Statistical Analysis* and Figure 2). (C) Mean saccade frequency (Hz) during the inhibition window and (D) during the rebound window, plotted across five successive flashes (flash order 1–5). Data are shown separately for foveal (blue) and parafoveal (orange) flash locations. While inhibition frequency remains stable across flashes, rebound frequency tends to diminish.

### Statistical modeling of inhibition frequency. Inhibition frequency is not modulated by flash repetition

We statistically tested these observations using linear mixed-effects models (LMEs) on parameters extracted within predefined time windows (see shaded rectangles in Figures 3, 4, and 5, and in the *Experimental Design and Statistical Analysis* section for a full description of the dependent measures). To quantify how the strength of saccadic inhibition varied across conditions, we modeled *Inhibition frequency*—defined as the mean saccade rate during the inhibitory dip—as a function of *Baseline frequency*, *Flash location*, *Flash order*, and their interaction, with random intercepts for each participant to account for repeated measures (Eq. 1). The model intercept was 1.36 Hz (p < 0.001), representing the average saccade frequency during inhibition under the reference condition (i.e., first flash, central location, and average baseline rate). This value reflects a typical reduction of ∼1.5 Hz compared to the pre-flash baseline (∼3 Hz, Figure 5), indicating robust oculomotor inhibition. A significant main effect of *Flash location* (β = 0.63, p < 0.001) revealed that inhibition was weaker when the flash occurred in the parafoveal visual field: saccade frequency during the inhibition period was higher at parafoveal compared to central locations (Figure 5A and B, blue vs. red lines and Figure 5C and D, all red dots above blue dots for all Flash orders). This supports the presence of spatial specificity in the strength of inhibition, consistent with known differences in foveal magnification across cortical and subcortical circuits (Cynader and Berman, 1972; Robinson, 1972; Benson et al., 2012; Hafed and Chen, 2016; Kupers et al., 2022; Fracasso et al., 2023).

Although baseline saccade frequency showed a declining trend across flash repetitions (Figure 5)—possibly reflecting fatigue or habituation — it did not significantly modulate inhibition frequency (β = 0.03, p = 0.545). Similarly, neither the main effect of Flash order (all p > 0.15) nor the Flash order by location interaction (all p > 0.07) reached significance. These results suggest that neither repetition nor individual baseline oculomotor state reliably influenced the depth of inhibition under these conditions. Finally, random effects revealed modest between-subject variability (SD = 0.51 Hz), indicating that average inhibition strength was relatively stable across participants. Overall, these findings support the interpretation that saccadic inhibition is a robust, reflex-like phenomenon—largely invariant to both recent stimulation history and individual differences in baseline saccade rates.

### Statistical modeling of inhibition latency. Inhibition onset scales with the baseline level of oculomotor activity

We next fit a linear mixed-effects model to predict Inhibition latency. The intercept was 121.93 ms (p < 0.001), consistent with classical estimates of saccadic inhibition timing under baseline conditions (Reingold and Stampe, 2002; Buonocore and McIntosh, 2008; Edelman and Xu, 2009; Bompas and Sumner, 2011), but with a slight delay, probably due to the free viewing nature of the task and the persistency of the visual transient which lasted 80 ms, probably widening the inhibitory dip. A significant negative effect of *Baseline frequency* (β = –5.23, p = 0.027) indicated that participants with higher pre-flash saccade rates tended to exhibit earlier inhibition. This suggests that the onset of inhibition scales with the initial level of oculomotor activity: participants with more active oculomotor states may generate stronger or more efficient inhibitory signals, reaching the suppression threshold faster. Conversely, lower baseline rates may reflect a less excitable oculomotor system requiring more time to initiate inhibition. A fixed-threshold mechanism, modulated by ongoing activity, could account for this variability in latency (Dorris and Munoz, 1998). Neither *Flash order* (all p > 0.20) nor *Flash location* (p = 0.80) yielded significant main effects, suggesting that these factors alone did not influence inhibition timing. However, the *Flash order-by-location* interaction for the fifth flash was significant (β = –23.89 ms, p = 0.048), suggesting a potential trend in which later flashes at parafoveal locations were associated with slightly earlier inhibition.

### Statistical modelling of inhibition frequency. Rebound responses decline with repetition

Unlike inhibition, rebound responses demonstrated clear habituation over time. Rebound frequency peaked approximately 250–300 ms after flash onset, followed by a gradual decline in amplitude across successive flashes (Figure 5A, B, C, and D). The intercept was 3.49 Hz (p < 0.001), reflecting the average saccade frequency during the rebound period under reference conditions. This value marks the typical post-inhibitory overshoot in eye movement activity (Reingold and Stampe, 2002; Buonocore and McIntosh, 2008; Edelman and Xu, 2009; Bompas and Sumner, 2011). A significant main effect of Flash order was observed in particular for the third and fifth repetition (Flash order3: β = –0.42, p = 0.04; Flash order₅: β = –0.68, p = 0.0016), indicating a progressive reduction in rebound frequency over time. This decline suggests that the oculomotor system became less responsive across repeated flashes—possibly reflecting habituation. A significant main effect of *Flash location* (β = –0.56, p = 0.008) confirmed that rebound strength was location-dependent, with reduced reactivation at parafoveal sites. Additionally, higher baseline saccade frequency predicted stronger rebound responses (β = 0.20, p = 0.003), suggesting that participants with more active oculomotor baselines not only inhibited earlier (see section above) but also re-engaged motor activity more robustly after suppression.

### Inhibition and rebound are dissociable processes

To determine whether rebound responses were simply a delayed consequence of stronger inhibition, we included *Inhibition frequency* as a predictor in the rebound model (see Eq. 2). *Inhibition frequency* had no effect (β = –0.15, p = 0.098), suggesting no predictive value. Furthermore, partial correlations between inhibition and rebound—controlling for baseline saccade frequency—were near zero across all conditions. These results support the notion that inhibition and rebound are functionally dissociable, reflecting distinct neural mechanisms: inhibition appears reflexive and sensory-driven, while rebound may reflect adaptive motor reactivation, potentially modulated by top-down influences.

Crucially, *Inhibition frequency* did not significantly predict rebound strength, indicating that these two components of the saccadic response—reflexive suppression and subsequent reactivation—can vary independently. The absence of a systematic relationship between inhibition depth and rebound amplitude argues against a simple compensatory mechanism and instead supports the view that these reflect distinct processes with separable sensitivities. Overall, these findings indicate that while rebound dynamics are sensitive to stimulus location, baseline activation, and stimulus repetition, they are not determined by the preceding degree of inhibition, further supporting their dissociation.

### Cumulative habituation across experimental blocks

To examine whether habituation effects accumulated over the course of the experiment, beyond within-trial repetition, we conducted an additional analysis assessing changes in oculomotor responses across experimental blocks. Trials were grouped into blocks of 150 trials, and saccadic inhibition and rebound magnitude were computed separately for each block. This analysis largely confirmed the results presented above, revealing a differential pattern across components. The magnitude of saccadic inhibition remained stable across blocks, indicating no progressive attenuation over extended exposure. In contrast, the rebound component showed a significant decrease across blocks, particularly between the first and fourth blocks (β = –0.56, p = 0.01), consistent with a cumulative habituation effect over the course of the experiment. Importantly, this block-level modulation did not interact with flash location (foveal vs. parafoveal) (all p > 0.1), suggesting that the long-term attenuation of rebound reflects a general adaptation process rather than a spatially specific effect. These findings indicate that while the immediate inhibitory response to visual transients remains robust even with extended exposure, the post-inhibitory rebound is progressively attenuated, supporting the interpretation that these two components exhibit distinct temporal dynamics.

### Rebound suppression shows experiment-specific modulation but no clear temporal decay

One possible explanation for the pattern of results observed in Experiment 1 is that the oculomotor system becomes less reactive over time during the free-viewing task, regardless of flash presentation. In this scenario, when probed, the sensory-driven inhibition remains intact, but the motor reprogramming component would gradually decrease over time. Crucially, this change would occur regardless of the number of flashes presented and would be simply related to the time dedicated to the task (*time-on-task*). This would alter the shape of the saccadic inhibition profile, yielding a stable inhibition level, a progressively reduced rebound across flash order, and a decrease in baseline frequency. To test this hypothesis, we conducted a second experiment, identical to Experiment 1, but where only the first or the fifth flash was presented. If the effect observed in Experiment 1 reflects habituation and time-on-task, motor rebound levels would be comparable between the first and fifth flash in Experiment 2. In contrast, if the effect were entirely due to time-on-task, differences between the first and fifth flash should still be observed under these conditions.

In Experiment 2, 30,767 saccades were retained after excluding detection errors. As in Experiment 1, saccade metrics conformed to known patterns of free viewing, including predominantly horizontal directions, a peak amplitude around 2.5 degrees of visual angle, a typical main sequence, and canonical temporal dynamics. Crucially, a reliable suppression of saccade frequency was observed around 130 ms after flash onset across participants, confirming the robustness of saccadic inhibition under naturalistic conditions.

To assess whether the rebound response varies with flash position, we compared *Rebound frequency* for the fifth flash between Experiments 1 and 2 (Figure 6). *Rebound frequency* differed significantly between experiments for the fifth flash (t = – 2.35, df = 38, p = 0.02). Moreover, while a clear reduction from the first to the fifth flash was observed in Experiment 1, no such difference was found in Experiment 2 (t = 1.5, df = 18, p = 0.15). This pattern is inconsistent with an explanation based on time-on-task effects, which would predict within-experiment differences even when only one flash was shown. Instead, the findings support a habituation account, specific to repeated flash stimulation, whereby repeated flash exposure progressively weakens the rebound phase of saccadic inhibition.

**Figure 6.**
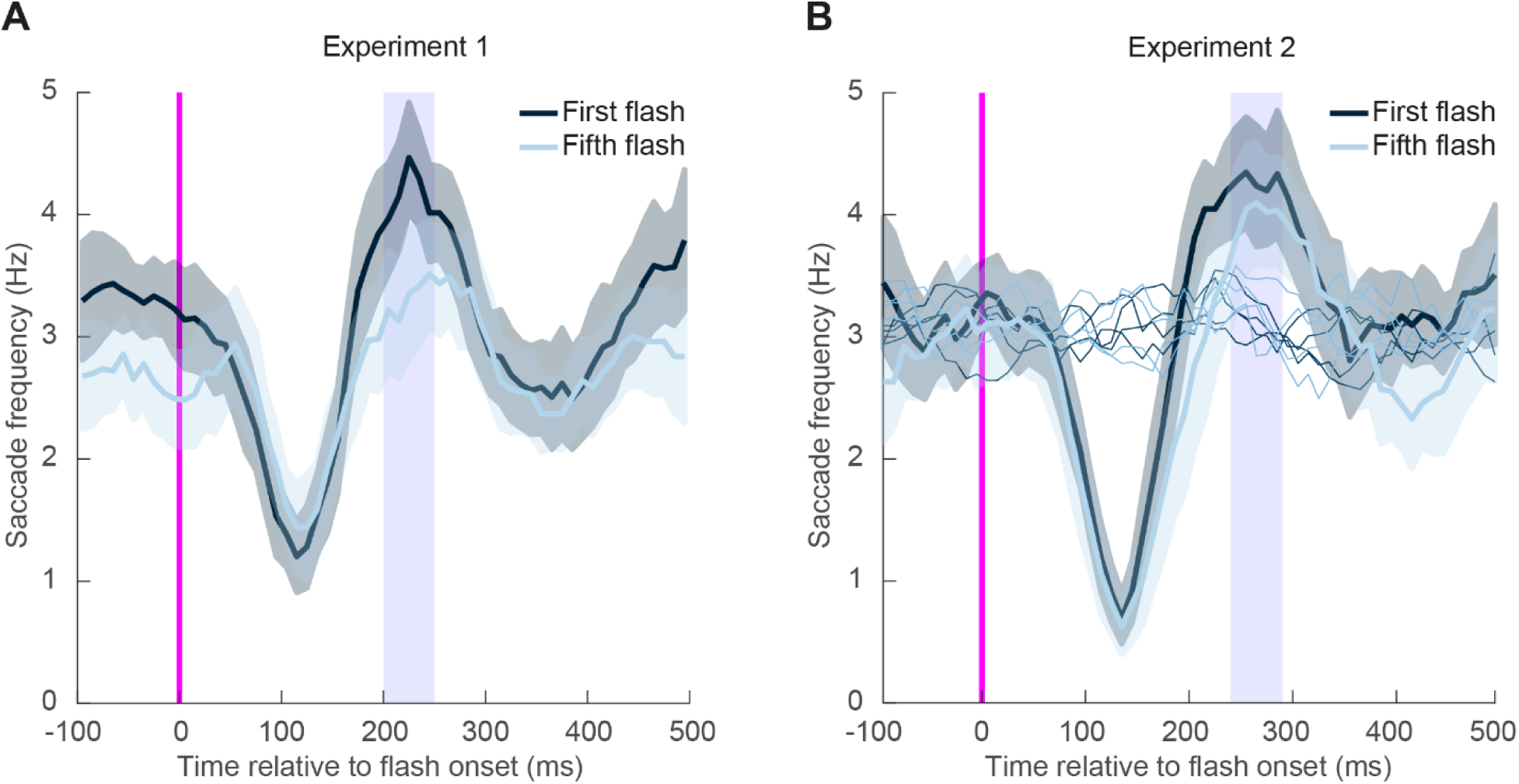
Saccadic inhibition profiles for first and fifth flashes in Experiment 1 and Experiment 2. (A) Experiment 1: SI profiles for the first (dark line) and fifth (light line) flash within a trial sequence. Inhibition depth remained stable, but rebound frequency was significantly reduced for the fifth flash. (B) Experiment 2: SI profiles when only the first or the fifth flash was presented per trial. No significant difference emerged between conditions, indicating that rebound suppression in Experiment 1 reflects repeated exposure rather than the flash’s temporal position. Shaded rectangles indicate analysis windows for baseline, inhibition, rebound, and post-rebound measures. In panel B, the thin lines refer to the “invisible” flashes that did not elicit any inhibition. As expected, when no stimuli were presented, the baseline level remained constant across the entire time window.

## Discussion

The present study investigated how saccadic inhibition evolves across repeated visual stimulation during naturalistic free viewing. Across two experiments, we found that the initial inhibition phase following a visual transient remained stable across repeated flashes, whereas the subsequent rebound—characterized by an increase in saccade rate above baseline—showed progressive attenuation (Experiment 1). Simple time-on-task effects (Shan and Edelman, 2023) could not account for the observed pattern: both inhibition and rebound magnitudes were comparable between the first and last transient when presented in isolation (Experiment 2). On the other hand, the rebound generated by the fifth flash in isolation was stronger than that generated by the fifth repeated flash (Experiment 1 vs. Experiment 2). Thus, we suggest that the decline in rebound in Experiment 1 reflects habituation induced by repeated sensory transients.

While the comparison between Experiments 1 and 2 relied on separate participant samples, the pattern of results strongly supports the interpretation that rebound attenuation is primarily driven by repeated exposure rather than by the nominal temporal position of the flash. Specifically, attenuation of the rebound emerged when flashes were repeated within a trial (Experiment 1), but was absent when only a single flash was presented (Experiment 2), despite identical timing parameters. It is important to note how this could also be tested with a within-subject manipulation of both paradigms. However, such a design could introduce carry-over or learning effects that might themselves alter habituation dynamics. Thus, while acknowledging the between-subject nature of the comparison, we therefore interpret the present findings as evidence that repeated exposure within a continuous viewing context is the critical factor underlying rebound attenuation.

### Sensory Inhibition and Motor rebound reflect distinct processes

The decoupling of inhibition and rebound is evidence that distinct mechanisms underlie the two phases of the oculomotor response to sudden visual changes. Specifically, the initial inhibition reflects a resilient, reflex-like suppression process (Reingold and Stampe, 2002; Buonocore and McIntosh, 2008; Edelman and Xu, 2009; Bompas and Sumner, 2011), whereas the rebound reflects an adaptive, experience-dependent modulation. Altogether, our findings demonstrate a functional dissociation between inhibition and rebound in oculomotor behavior during continuous free viewing.

Our results extend previous work on SI adaptation (Shan and Edelman, 2023) by showing that the inhibitory phase is adapted only within a specific time window. Unlike the rapid attenuation of the inhibition reported at short ISIs (∼0.2 s), the absence of SI decline at longer flash intervals (∼0.7–1.2 s) suggests that low-level sensory adaptation recovers rapidly over time, in the order of a few hundred milliseconds (Wark et al., 2007). This mechanism may be particularly relevant under free-viewing conditions, in which inhibition remains stable to support the reinstatement of saccades, which are often redirected toward new visual targets (Buonocore et al., 2017a). The present findings should not be interpreted as evidence for anatomically independent neural processes underlying the inhibitory trough and the rebound. Instead, the data support a functional dissociation between two temporally separable components of the oculomotor response to visual transients. These components may reflect partially independent processes with distinct adaptation profiles, or they may arise from different operating states within a common sensorimotor circuit. Resolving this issue will require converging evidence from paradigms that combine behavioral measures with neurophysiological or neuroimaging techniques, as well as computational modeling capable of testing whether a single dynamical system can account for both inhibition and rebound.

### Habituation in the motor system

This pattern of stable inhibition, contrasted with weakening motor output, resembles habituation in other motor systems, in which early sensory gating is preserved while subsequent motor responses decline. For example, in the vestibulo-ocular reflex (VOR)—which stabilizes vision by producing eye movements opposite to head movements—the basic sensory pathway that detects head motion remains intact. However, when the system is exposed to repeated or prolonged stimulation, the motor response gradually weakens. This reduction in “motor gain” means that the eyes no longer fully counteract the head movement, resulting in progressively smaller compensatory eye movements (Lisberger et al., 1994). In another sensory domain, the acoustic startle reflex—the rapid whole-body flinch triggered by sudden loud sounds—illustrates a similar pattern (Yeomans and Frankland, 1995). The initial inhibitory process that suppresses background activity remains consistently engaged, ensuring that the sound is detected. However, when the stimulus is repeated, the strength of the motor response (the actual body movement) gradually diminishes. In other words, the system continues to register the sound but produces a progressively smaller behavioral response over time (Davis, 1984; Koch, 1999; Zheng and Schmid, 2023).

At the neuronal level, these behavioral effects correspond to distinct sensory and motor dynamics. In the superior colliculus (SC), responses to repeated visual transients decrease despite stable sensory detection (Boehnke et al., 2011). In brainstem circuits underlying the startle reflex, neurons exhibit early inhibition of auditory input, along with diminished motor output with repetition (Davis, 1984; Geyer and Braff, 1987). Across these systems, inhibitory gating mechanisms remain intact to preserve responsiveness, whereas motor re-engagement is selectively attenuated. By analogy, the dissociation we observed between stable inhibition and weakening rebound in SI likely reflects the same principle: robust sensory suppression coupled with an adaptive, experience-dependent reduction in motor reprogramming under repeated stimulation.

### Neural mechanisms

While the current manuscript centers on behavioral (oculomotor) data, we can speculate on the most likely neural mechanism mediating the observed results. The persistence of SI across repetitions is consistent with existing models proposing subcortical circuits that gate saccade execution. Within the oculomotor system, omnipause neurons (OPNs) in the pontine reticular formation exert tonic inhibition over saccade-generating burst neurons (Fuchs and Luschei, 1972; Luschei and Fuchs, 1972; Keller, 1974; Büttner-Ennever and Büttner, 1978; Evinger et al., 1982; Büttner-Ennever et al., 1988; Büttner-Ennever et al., 1999; Takahashi et al., 2022), maintain tonic firing during fixation and pausing just before saccade initiation. Recently, we have suggested that OPNs may receive direct sensory input, allowing rapid responses to exogenous stimuli and raising the threshold for saccade initiation (Buonocore and Hafed, 2023). This would explain the rapid onset of SI following a new stimulus. Increases in OPN activity following visual flashes might reflect a hard-wired gating mechanism that operates independently of behavioral relevance. Supporting this, the timing and magnitude of SI in our data remained invariant even under repeated stimulation, consistent with evidence that either direct reactivation of OPNs or visual bursts in the SC via projections to OPNs can rapidly trigger suppression of saccade generation.

In contrast, the rebound may rely on mechanisms that disengage OPN activity and re-excite burst neurons. Repeated stimulation may reduce the efficiency of this disengagement, leading to weaker reactivation of saccade commands. Such selective habituation echoes findings in the SC (Boehnke et al., 2011; Basso and May, 2017) and frontal eye fields (FEF) (Bruce and Goldberg, 1985; Mayo and Sommer, 2008), where responses to repeated transients diminish despite preserved detection.

These findings imply a recalibration of the balance between inhibitory stability and reorienting flexibility within brainstem circuits. Our work provides a systems-level perspective on how sensorimotor loops evolve during active vision. SI thus emerges from dynamic interactions between SC, FEF, and OPNs. While SC and FEF provide excitatory signals for orienting, OPNs contribute an inhibitory signal that prevents premature eye movements. By ensuring that saccades are executed only after sufficient sensory processing, this interaction adapts visual behavior to environmental demands. Variations in attentional state, relevance, or motivational salience may gradually disengage the system from reflexive orienting responses.

Interestingly, the temporal structure of the rebound included dynamics consistent with rhythmicity in oculomotor control (see Figures 3A and 3B, and 4A and 4B). Recurrent fluctuations in saccade likelihood following inhibition may be shaped by beta-or alpha-band oscillations in frontoparietal areas, which have been linked to saccadic timing and attentional cycles (VanRullen, 2016; Gaillard et al., 2020). This rhythmic post-inhibitory signature may support a framework in which the brain evaluates salience in periodic windows, reconfiguring sensorimotor coupling accordingly (Helfrich et al., 2018; Fiebelkorn and Kastner, 2019).

## Conclusion

Together, these findings reveal a selective habituation of motor re-engagement in SI, whereas sensory-driven inhibition remains robust. This dissociation shows that the oculomotor system recalibrates to repeated stimulation by preserving inhibitory fidelity while attenuating rebound, thereby filtering out irrelevant events. More broadly, SI offers a window into how sensory and motor components of eye-movement control adapt independently, with implications for both basic mechanisms and clinical applications. Functionally, a system that maintains transient suppression yet reduces redundant reprogramming may balance sensory prioritization with motor efficiency. In cluttered or predictable environments, minimizing rebound to repeated flashes could optimize exploration by limiting distraction and conserving resources.

This separation also holds clinical relevance. Disorders such as Parkinson’s disease and ADHD are characterized by deficits in oculomotor inhibition and variability in response timing (Antoniades and Spering, 2024). In Parkinson’s disease (PD), in addition to fatigue in movement control (Di Vico et al., 2021), oculomotor abnormalities also emerge early in the disease course. Patients typically exhibit saccadic hypometria, prolonged latencies, and reduced peak velocities (Pretegiani and Optican, 2017). Performance on tasks probing reflexive oculomotor control—such as visually guided prosaccades—is often relatively preserved.

In contrast, antisaccade tasks consistently reveal impaired voluntary inhibition and strategy reconfiguration, with increased error rates and longer latencies, including at early drug-naïve stages (Antoniades et al., 2015). Against this backdrop, our results point to a clinically relevant dissociation that is translatable to clinical implementation for PD assessment: the SI dip (transient, stimulus-driven suppression of saccades) appears preserved, whereas the motor rebound is probably delayed and/or reduced in magnitude and time relative to controls—consistent with a framework in which sensory-driven inhibitory gating remains intact while motor re-engagement is selectively compromised (Fooken et al., 2022; Buonocore and Hafed, 2023). This pattern aligns with evidence that some oculomotor behaviors relying on superior colliculus–brainstem pathways are spared in PD, whereas behaviors requiring greater engagement of the fronto–basal ganglia are disproportionately affected. The ability to quantify the stability of SI and the adaptability of rebound may yield sensitive markers for early detection of dysfunctions affecting automatic and voluntary control systems.

Future work should investigate neural signatures of this dissociation using EEG or fMRI and assess whether rebound adaptation generalizes across modalities and tasks. By exploring the interaction between SI and sensory adaptation, we aim to uncover the principles that regulate visual exploration. This knowledge could inform both theoretical models of oculomotor control and translational applications in perception and clinical neuroscience.

## Acknowledgments

A.B., C.C., G.C., and G.V are supported by a grant from the Italian Minister of Research and University (PRIN 2022_PNRR-P2022ST78T). C.C. and G.C. were supported by #NEXTGENERATIONEU (NGEU) and funded by the Ministry of University and Research (MUR), National Recovery and Resilience Plan (NRRP), project MNESYS (PE0000006) – A Multiscale integrated approach to the study of the nervous system in health and disease (DN. 1553 11.10.2022). A.F. was supported by a grant from the Biotechnology and Biology Research Council (BBSRC, grant number: BB/S006605/1) and the Bial Foundation (Bial Foundation; Grant id: A-29315, number: 203/2020, grant edition: G-15516). The authors used AI-assisted technology for grammar checking of the manuscript text.

